# Multilocus Species Trees Show the Recent Adaptive Radiation of the Mimetic *Heliconius* Butterflies

**DOI:** 10.1101/003749

**Authors:** Krzysztof M. Kozak, Niklas Wahlberg, Andrew Neild, Kanchon K. Dasmahapatra, James Mallet, Chris D. Jiggins

## Abstract

Müllerian mimicry among Neotropical Heliconiini butterflies is an excellent example of natural selection, and is associated with the diversification of a large continental-scale radiation. Some of the processes driving the evolution of mimicry rings are likely to generate incongruent phylogenetic signals across the assemblage, and thus pose a challenge for systematics. We use a dataset of 22 mitochondrial and nuclear markers from 92% of species in the tribe to re-examine the phylogeny of Heliconiini with both supermatrix and multi-species coalescent approaches, characterise the patterns of conflicting signal and compare the performance of various methodological approaches to reflect the heterogeneity across the data. Despite the large extent of reticulate signal and strong conflict between markers, nearly identical topologies are consistently recovered by most of the analyses, although the supermatrix approach fails to reflect the underlying variation in the history of individual loci. The first comprehensive, time-calibrated phylogeny of this group is used to test the hypotheses of a diversification rate increase driven by the dramatic environmental changes in the Amazonia over the past 23 million years, or changes caused by diversity-dependent effects on the rate of diversification. We find that the tribe Heliconiini had doubled its rate of speciation around 11 Ma and that the presently most speciose genus *Heliconius* started diversifying rapidly at 10 Ma, likely in response to the recent drastic changes in topography of the region. Our study provides comprehensive evidence for a rapid adaptive radiation among an important insect radiation in the most biodiverse region of the planet.

---

Visual mimicry provides an excellent system in which to study the origins of biodiversity, as the targets of selection are clearly identifiable and the role of natural selection in promoting adaptation and ultimately speciation can be directly observed (Bates 1863; Benson 1972; Mallet and Barton 1989; Sherratt 2008; Pfennig 2012). Studies of mimetic assemblages have been instrumental in explaining many biological phenomena, ranging from reproductive isolation (McMillan et al. 1997; Jiggins et al. 2001; Merrill et al. 2012) to genetics of adaptation (Benitez-Vieyra et al. 2007; Baxter et al. 2009; Jones et al. 2012), and including prominent examples from vertebrates (Brodie III and Brodie Jr 2004, Wright 2011), arthropods (Ceccarelli and Crozier 2007, Janzen et al. 2010, Hines and Williams 2012), plants (Benitez-Vieyra et al. 2007), and microbes (Elde and Malik 2009).

Testing hypotheses regarding the evolution of mimicry depends heavily on our knowledge of systematic relationships between the participating taxa, particularly where mimics are closely related. Phylogenetic comparative approaches have been used to address questions such as concerted coevolution of Müllerian co-mimics in catfish (Wright 2011), advergence due to directional selection in Batesian mimicry of spiders (Ceccarelli and Crozier 2007) and selection on imperfect mimics among the hoverflies (Penney et al. 2012). Unfortunately, both hybridisation driven by strong selection on adaptive loci and rapid radiation leading to incomplete lineage sorting are likely to be common in mimetic systems (Savage and Mullen 2009; Kubatko and Meng 2010; Kunte et al. 2011; Zhang et al. 2013) and can significantly interfere with the estimation of phylogenies (Maddison and Knowles 2006; Linnen and Farrell 2008; Edwards 2009; Anderson et al. 2012).

We use the Neotropical butterfly tribe Heliconiini (Nymphalidae: Heliconiinae) to explore the problem of phylogeny estimation in a Müllerian mimetic group and investigate the link between the dynamics of speciation and macroevolutionary factors. Heliconiini are arguably the most thoroughly-researched example of microevolution in the Neotropics, the most biodiverse region of the world (Hoorn et al. 2010). They comprise the genus *Heliconius* and nine smaller genera, providing a spectacular example of a radiation where speciation is frequently caused by mimicry of aposematic patterns directed against avian predators (Jiggins et al. 2001; Arias et al. 2008; Merrill et al. 2012). The system has provided an excellent example for the study of convergence from both genomic and organismal perspective (e. g. Duenez-Guzman et al. 2009; Hines et al. 2011; Bybee et al. 2012; Heliconius Genome Consortium 2012; Martin et al. 2012, 2013; Pardo-Díaz et al. 2012; Jones et al. 2013; Nadeau et al. 2013; Supple et al. 2013). The clade presents an opportunity for comparative studies of many unique traits, including larval and imaginal sociality (Beltrán et al. 2007), pupal mating (Estrada et al. 2011), pollen feeding (Cardoso and Gilbert 2013), or close association with varying numbers of Passifloraceae host plants (Brown 1981; Merrill et al. 2013). The ongoing proliferation of comparative molecular and genomic studies on the diversity of genes regulating vision (Pohl et al. 2009), chemosensation (Briscoe et al. 2013) and cyanogenesis (Chauhan et al. 2013) makes a stable molecular phylogeny especially desirable.

An unusual feature of *Heliconius* is the prevalence and importance of gene flow and hybridisation, leading to a controversy over the validity of traditional species concepts in the clade (Beltrán et al. 2002; Mallet et al. 2007). At least 26% of all species of Heliconiini occasionally produce hybrids in the wild (Mallet et al. 2007), and *H. heurippa* has resulted from homoploid hybrid speciation from parental forms diverged millions of years ago (Mavárez et al. 2006; Jiggins et al. 2008; Salazar et al. 2010). Genome sequencing has shown that mimetic diversity in the *H. melpomene* and silvaniform clades is also explained by adaptive introgression of genes regulating the aposematic wing patterns (Heliconius Genome Consortium 2012; Pardo-Díaz et al. 2012), and seemingly neutral gene flow is widespread, influencing as much as 40% of the genome between *H. melpomene* and *H. cydno* (Martin et al. 2013).

A large body of recent work has been devoted to the issue of incongruence between the species tree (the true speciation history) and the gene trees evolving within (Anderson et al. 2012; Cutter 2013). The traditional approach of concatenating the total genetic evidence into a supermatrix in order to obtain a global estimate of the predominant phylogenetic signal and hidden support (Gatesy and Baker 2005), without consideration for the heterogeneity of individual partitions, has been to some extent superseded by multispecies coalescent (MSC) techniques (reviewed in: Edwards 2009; Knowles and Kubatko 2010; Anderson et al. 2012; Cutter 2013; Leaché et al. 2013). The majority of the new MSC algorithms are intended to model at least some of the sources of heterogeneity between different markers, most frequently focusing on the problem of incomplete lineage sorting (e.g. Maddison and Knowles 2006; Heled and Drummond 2010), sometimes addressing hybridisation (e.g. Gerard et al. 2011; Yu et al. 2011), and in at least one case modelling discordance without specifying its potential source (Larget et al. 2010). Although the supermatrix approach remains popular and serves as an effective approximation of the species diversification history in most cases, its ability to properly assess the degree of statistical support for phylogenies has been brought into question and contrasted with the potential of the MSC techniques to assess confidence more realistically (Edwards 2009; Knowles and Kubatko 2010). Heliconiini are an especially interesting subject for a systematic study, where the purported robustness of MSC tools to gene flow and other sources of incongruence can be tested with real data and used to evaluate support in the light of underlying heterogeneity of the phylogenetic signal.

Heliconiini systematics has a long history, starting with early morphological work (Emsley 1965; Brown 1981; Penz 1999), through allozymes (Turner et al. 1979) and a combination of morphological and RNA-restriction data (Lee et al. 1992), to studies based on the sequences of mitochondrial and nuclear markers (Brower 1994b; Brower and Egan 1997; Beltrán et al. 2002, 2007; Cuthill and Charleston 2012). The most comprehensive study to date by Beltrán et al. (2007) attempted to address some of the difficulties by incorporating many taxa (38 of 46 *Heliconius*, 59 of 78 Heliconiini), considering two individuals of most species, and sequencing two mitochondrial (*CoI/II*, *16S*) and four nuclear markers (*EF1α*, *Wg*, *Ap*, *Dpp*). However, this dataset is still potentially inadequate to address the challenges posed by Heliconiini systematics, as three of the loci (*16S*, *Ap*, *Dpp*) were only sequenced for 12 representative species. Among the other three markers, 65% of the variable sites resided in the fast-evolving mitochondrial partition *CoI/II*, raising the possibility that the inferred relations are largely driven by the historical signal of the matriline. The proposed relationships between the basal genera of the tribe were poorly supported and the relationships between *Heliconius*, *Eueides* and the other eight genera was not resolved, as might be expected if most of the data comes from a fast-evolving partition. Importantly, the individual data partitions were analysed in concatenation, as the debate on the species tree-gene tree incongruence was in its early stages at the time. Despite these shortcomings, Beltrán and colleagues confirmed that the morphological and behavioural characteristics of the major clades do not correspond directly to their evolutionary history and that the traditionally recognised genera *Laparus* and *Neruda* are most likely nested within the crown genus *Heliconius*.

The importance of Müllerian mimicry as a driver of individual speciation events has been well-established (Mallet et al. 1998; Mallet and Joron 1999; Jiggins 2008) whereas the macroevolutionary processes governing the evolution of the group have been largely neglected in empirical studies (but see Etienne et al. 2012; Rosser et al. 2012). Precise understanding of the evolution of the Müllerian mimicry rings as well as associated processes such as hybridisation, requires knowledge of the relative timing of the divergence events and motivated the most widely-cited study of the molecular clock in Arthropoda (Brower 1994a). Mallet et al. (2007) used a relaxed clock procedure to adjust the original branch lengths based on the *CoI/II* alignment. However, some of the relations were left unresolved, and the sequences were not partitioned into fast and slow-evolving codon positions, which can result in inflated age estimates (Brandley et al. 2011).

Most importantly, the first dated phylogeny of Heliconiini is not calibrated to an absolute standard, making it impossible to make inferences about the relation of the diversification process and the contemporaneous geological and climatic events. The cumulative results of over 200 systematic studies demonstrate that most South American tropical clades have experienced periods of significantly elevated net diversification rate in response to Andean orogenesis, alterations in the hydrology and sediment dynamics of the present-day Amazon Basin, as well as local and global climatic changes (Hoorn et al. 2010; Turchetto-Zolet et al. 2013). These processes can result in allopatry of incipient lineages, or in creation of new ecological niches and change to the species-level carrying capacity of the environment leading to ecological speciation (Brumfield and Edwards 2007; Hoorn et al. 2010). A recent comparative study suggests that the majority of *Heliconius* lineages originated in the North-Eastern Andes and spread to other parts of the continent (Rosser et al. 2012). Three periods stand out as potentially critical in the history of Heliconiini as the times when orogenic, hydrologic and climatic events would have created new habitats and altitudinal gradients. The first phase of the intense Andean uplift, taking place 23 Ma, was also the start of the great marine incursion into the North of the continent, separating the incipient mountain range from the Eastern plateaus (Gregory-Wodzicki 2000; Solomon et al. 2008; Hoorn et al. 2010). 12.4 Ma marked the end of the Mid-Miocene Climatic Optimum and the onset of global cooling (Lewis et al. 2007), which likely led to the reduction and separation of the rainforest areas by savannas unsuitable for many Heliconiini (Jaramillo et al. 2011). Coincident was the second stage of rapid Andean orogeny 12 Ma, strongly changing the elevation gradients in the Central and Eastern sectors (Gregory-Wodzicki 2000), and followed shortly by the entrenchment of the Amazon in its modern course 10 Ma, and a gradual expansion of the rainforest in place of the retreating wetlands during the next 3 Myr (Hall and Harvey 2002, Figueiredo 2009, Hoorn 2010). Finally, the Eastern Colombian Cordillera uplifted in the last burst 4.5 Ma and the Isthmus of Panama connected at least 3 Ma, possibly changing the patterns of isolation between various populations (Hill et al. 2013). Hence we can hypothesise that the diversification rate of Heliconiini increased during the periods of intensive Andean uplift around 23, 12 and 4.5 Ma.

## Aims of the Study

Here we aim to resolve the species tree of the Heliconiini radiation and generate a dataset including virtually all of the currently valid species in the tribe, sampling intraspecific diversity across the range of many of the species. We include information from 20 nuclear and two mitochondrial markers, as well as the whole mitochondrial sequence of select species. The exonic markers comprise both autosomal and Z-linked loci, chosen to represent a sampling of chromosomes and a range of evolutionary rates, from fast-evolving mitochondrial genes (e.g. *CoI/II*) to more slow evolving nuclear loci (e.g. *Wg*). We apply a wide range of phylogenetic methods to reconstruct the species tree, including supermatrix, coalescent and network approaches, which allow us to assess the strength of the underlying signal of speciation. The power of our combined approach is harnessed to detect the incongruence between markers, likely to arise from the complex population-level processes acting in a radiation of Müllerian co-mimics. We elucidate the importance of marker heterogeneity for the final assessment of systematic relationships, while realistically estimating the support values for our chosen topology. A robust estimate of the relationships between 92% of the species and multiple outgroups makes it possible to date the time of individual divergence events with confidence, providing input for an analysis of diversification dynamics. The calibrated chronogram is used to test our hypothesis of diversification rate increase at times of dramatic environmental changes in the Andes and Amazonia, as well as the previous suggestions of diversity-dependent cladogenesis in Heliconiini (Fordyce 2010; Etienne et al. 2012). We thus present a comprehensive study of macroevolutionary dynamics in a mimetic system that has been studied intensively at the microevolutionary level.

## Materials and Methods

### Taxon Sampling

We sampled 180 individuals, including 71 of the 78 species in all genera of Heliconiini and 11 outgroup species. The specimens came primarily from our collection at the University of Cambridge, with additional specimens shared by museums and private collectors (Online Appendix 1). We included five outgroup species from the sister tribe Acraeini (Wahlberg et al. 2009) and three from the related genus *Cethosia*. The diverse analyses used in this paper require different sampling designs and the demands of all the techniques cannot be easily accommodated in a single dataset. For example, the network analysis based on nucleotide distance produced much better supported and resolved trees when the 95% or more incomplete data from historical specimens were not used, whereas the various multi-species coalescent techniques required the use of at least two individuals per species and had to be based only on taxa with intraspecific sampling (Fulton and Strobeck 2009; Heled and Drummond 2010). Thus we distinguish four datasets. The <u>complete</u> data matrix includes all the data. The <u>core</u> dataset excludes 14 individuals represented solely by short DNA fragments from historical specimens. The <u>single-individual</u> dataset includes both modern and historical specimens, but with only the single best-sequenced individual per taxon. Finally, the <u>*BEAST matrix</u> contains only the 17 species of *Eueides* and *Heliconius* with extensive sampling of multiple representatives of each species.

### DNA Sequencing

We used 20 nuclear and two mitochondrial loci as markers (Table 1; Online Appendix 1), exceeding the number of loci considered sufficient for most phylogenetic problems (Gatesy et al. 2007). The selection includes the three classic molecular markers for Lepidoptera (*CoI/II*, *EF1α*, *Wg*), two markers proposed by Beltrán et al. (2007)(*16S*, *Dpp*), eight new universal markers proposed by Wahlberg and Wheat (2008) (*ArgK*, *Cad*, *Cmdh*, *Ddc*, *Idh*, *Gapdh*, *Rps2*, *Rps5*) and nine highly variable loci identified by Salazar et al. (2010) (*Aact*, *Cat*, *GlyRS*, *Hcl*, *Hsp40*, *Lm*, *Tada3*, *Trh*, *Vas*). Additional *Heliconius*-specific primers were designed for *Cmdh*, *Gapdh* and *Idh*. Details of the primers and PCR cycles are listed in the Online Appendix 2. For most species, sequences of the three basic markers for multiple individuals were already published (Brower 1997; Beltrán et al. 2007; Wahlberg et al. 2009; Salazar et al. 2010), and data for 26 individuals came exclusively from GenBank (Online Appendix 1).

**Table 1.**
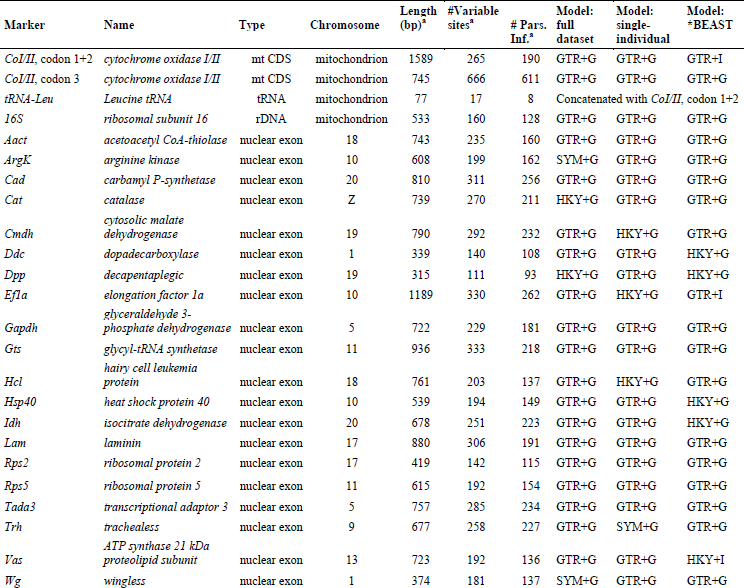
Position in the genome, variability statistics and models of sequence evolution for two mitochondrial and 20 nuclear markers used in the study. Statistics are reported for the core alignment including outgroups.

We generated Sanger sequences for 103 specimens (Online Appendix 1). DNA was isolated from approximately 50 μg of thorax tissue using the DNeasy Blood & Tissue kit (Qiagen, Manchester, UK). PCR was carried out in a total volume of 20 μl, containing 1× Qiagen Taq buffer (Manchester, UK), 2.5 mM MgCl_2_, 0.5 μM of each primer, 0.2 mM dNTPs, 1 unit bovine serum albumin, 0.5 unit Qiagen Taq-Polymerase and 1 μl of the DNA extract. The following program was executed on a G-Storm cycler (Somerton, UK): denaturation 5 minutes at 94°C; 35 cycles of 30 seconds at 94°C, 30 seconds at the annealing temperature and 90 seconds at 72°C; final extension for 10 minutes at 72°C. The results were visualised by electrophoresis in 1.5% agarose gel stained with 1% ethidium bromide. PCR products were cleaned using the ExoSAP-IT system (USB, Cleveland, Ohio): 60 minutes at 37°C; 20 minutes at 80°C. We used gel purification with the Nucleo Spin Extract II kit (Macherey-Nagel, Dueren, Germany) as needed. Sanger sequencing reaction was carried out with the BigDye Terminator v. 3.1 (AB, Foster City, California: 2 minutes at 94°C; 25 cycles of 10 seconds at 94°C, five seconds at 50°C and four minutes at 60°C. The products were sequenced with the ABI 3730×l DNA Analyzer at the Sequencing Facility, Department of Biochemistry, University of Cambridge. We manually inspected the traces in CodonCode v. 4.0.4 using PHRED for quality assessment (CodonCode Corporation 2012).

At the time of our Sanger sequencing effort, whole genome data generated for other studies became available from 57 individuals in 27 common species (Heliconius Genome Consortium 2012; Briscoe et al. 2013; Supple et al. 2013; Dasmahapatra and Mallet, unpublished) (Online Appendix 1). 100 base pair reads were generated using the Illumina Genome Analyzer II and HiSeq 2000 platforms with insert size of 300-400 bp. We performed *de novo* assembly of the short reads in the program Abyss v. 1.3 (Simpson et al. 2009). Based on previous studies (Salzberg et al. 2012; Briscoe et al. 2013) and preliminary results (S. Baxter, *pers. comm.*), we chose k-mer length of 31, minimum number of pairs n=5 and minimum mean coverage c=2 as optimal settings. The 20 nuclear markers were mined from the assemblies by megaBLAST (Camacho et al. 2009) in Geneious v. 5.5.1 (Biomatters Ltd 2012) using reference sequences from the *Heliconius melpomene* genome. The quality of the recovered sequences was assessed by alignment to previously generated amplicon sequences of the same loci from the same individuals.

Mitochondrial sequences could not be recovered by *de novo* methods, presumably because the large number of reads from the highly abundant mitochondrial DNA contained a high enough number of erroneous sequences to interfere with the assembly. We reconstructed whole mitochondrial genomes of 27 species by mapping to the *Heliconius melpomene* reference (The *Heliconius* Genome Consortium, 2012), using the default settings in the Genomics Workbench v. 5.5.1 (CLCBio 2012). This data was analysed separately from the 21 locus mixed nuclear-mitochondrial alignment. All the original sequences were deposited in GenBank (Accession numbers – data in submission; Online Appendix 1).

### DNA Sequencing: Historical Specimens

Short fragments of *CoI/II* and *EF1α* were sequenced from historical specimens up to 150 years of age, obtained from museum and private collections (Online Appendix 1) and processed in a vertebrate genetics laboratory to reduce the risk of contamination. Instruments and surfaces were cleaned with 5% bleach and irradiated with UV for 30 minutes prior to use. One to two legs were washed in water, immersed in liquid nitrogen in a test tube for 30 seconds and ground up, followed by an extraction into 20 μl of buffer using the QIAmp DNA Micro Kit (Qiagen, Manchester, UK). We treated every fifth extraction and every fifth PCR as a negative control with no tissue or DNA extract. PCR reactions were carried out in a 20 μl volume using 1 unit of Platinum HiFi Taq Polymerase (Invitrogen, London, UK) and 1× buffer, 2.5 mM MgCl_2_, 0.5 μM of each primer, 0.2 mM dNTPs, 1 unit bovine serum albumin, sterilised DNAse-free water and 1-5 μl of the DNA extract depending on concentration. In order to accommodate shearing of DNA with time, we designed and applied PCR primers spanning short fragments of 200-300 bp (Online Appendix 2). We carried out amplification, product clean up and sequencing as above, partially accounting for possible cross-contamination by blasting the results against GenBank.

### Alignment and Gene Tree Estimation

Alignments for each locus were generated in CodonCode v. 3 to account for inverted and complemented sequences, and improved using MUSCLE v. 3.8 (Edgar 2004). We visualised the alignments of the coding loci (all except the mitochondrial *16S* and the *tRNA-Leu* fragment in the *CoI/II* sequence) in Mesquite v. 2.75 (Maddison and Maddison 2011) and checked translated sequences for stop codons indicating errors. The whole mitochondrial sequences were aligned to the *Acraea issoria* and *Heliconius melpomene* references (*Heliconius* Genome Consortium 2012) using the G-INS-i algorithm in MAFFT (Katoh 2002). The number of variable and parsimony informative sites was estimated for each locus in PAUP* v. 4 (Swofford 2002). Models of sequence evolution implemented in MrBayes (Ronquist and Huelsenbeck 2003) and BEAST (Drummond and Rambaut 2007) were selected in MrModelTest v. 2.3 (Posada and Crandall 1998; Nylander 2004) based on the Akaike’s AIC (Akaike 1974). Xia’s test in DAMBE v. 4.0 (Xia and Xie 2001) demonstrated saturation in the third codon position of *CoI/II*. Protein-coding mitochondrial markers often display high saturation at the third codon position, potentially leading to incorrect parametrisation of substitution rate models (Brandley et al. 2011). To account for this effect, in all the subsequent analyses we treated the third codon position of the fast-evolving *CoI/II* locus as a separate partition. The Leucine tRNA (*tRNA-Leu*) fragment occurring in the middle of *CoI/II* displays very low variability and thus was included in one partition with the slower evolving first and second codon positions. Individual gene trees were estimated in MrBayes v. 3.1, using four runs of one chain, 10 million MCMC cycles sampled every 1000 cycles, and 2.5 million cycles discarded as burnin based on the convergence diagnostic. The mitochondrial genes were concatenated due to their shared history, but treated as separate partitions with distinct models. All trees were visualised with FigTree v. 1.4 (Rambaut 2009).

### Detection of Conflicting Signals

We investigated the cyto-nuclear discordance and other conflicts in the phylogenetic signal with several methods. To illustrate the global reticulate signal in the data, a NeighborNet network was built in the program SplitsTree v. 4 (Kloepper and Huson 2008) with the pairwise distances calculated under the F84 correction for multiple hits. We reduced the dataset to a single best-sequenced individual per species in order to exclude the reticulations resulting from the expected recombination within species. Next, the topological disparity among individual loci was illustrated using Multi-Dimensional Scaling of pairwise Robinson-Foulds distance (Robinson and Foulds 1981) between the gene trees, as estimated by TreeSetViz v. 1.0 (Hillis et al. 2005) in Mesquite. Calculation of the RF required trimming the trees to the minimal set of 54 shared taxa from 27 species, using the R package APE (Paradis et al. 2004; R Development Core Team 2008). Finally, we investigated if topologies and branch lengths of the individual loci are consistent enough to be concatenated, by means of a hierarchical likelihood ratio test in Concaterpillar v. 1.5 (Leigh et al. 2008).

### Supermatrix Phylogenetics

We created a supermatrix of the 20 nuclear and 2 mitochondrial markers in Mesquite and estimated the Maximum Likelihood (ML) phylogeny under the GTRGAMMA model in RAxML v. 7.0.4, with 1000 bootstrap replicates under the GTRCAT approximation (Stamatakis 2006). To explicitly test the likelihood of various hypotheses for Heliconiini phylogeny, several alternative topologies were created in Mesquite, representing previously identified groupings, as well as suggested placements of the enigmatic genera *Cethosia*, *Laparus* and *Neruda* (Beltrán et al. 2007, Brower 1994, Brower & Egan 1997, Mallet et al. 2007, Penz 1999, Penz & Peggie 2003). We then re-estimated the ML tree under the GTRGAMMA model using each topology as a constraint. The likelihood scores of the original and alternative trees were compared using the Shimodaira-Hasegawa test (Shimodaira and Hasegawa 1989) and the Expected Likelihood Weights based on 1000 bootstrap replicates (Strimmer and Rambaut 2002). A separate phylogeny was generated for the unpartitioned whole mitochondrial alignment in RAxML under the GTRGAMMA model with 1000 bootstrap replicates.

We estimated a calibrated Bayesian phylogeny using the program BEAST v. 1.7.5 (Drummond et al. 2006). In order to avoid incorrect estimates of the substitution rate parameters resulting from the inclusion of multiple samples per species, this analysis was based on a pruned alignment with one individual per species. The only exception is the inclusion of 3 races of *H. melpomene* and 2 races of *H. erato*, where deep geographical divergences are found (Quek et al. 2010). The Vagrantini sequences were not used, as BEAST can estimate the placement of the root without an outgroup (Drummond et al. 2006). Thus the analysis included 78 taxa and the substitution rate models were re-estimated appropriately (Table 1). We linked the topology, but modelled an uncorrelated lognormal clock and the substitution rate separately for each locus (Drummond et al. 2006). Substitution rates were drawn from the overdispersed gamma distribution prior with shape parameter k=0.001, scale parameter theta=1 and starting value 0.001 for nuclear genes, and k=0.01 for the faster-evolving mitochondrial loci (van Velzen 2013). We used a Birth-Death tree prior and empirical base frequencies to limit the computation time for the heavily parametrised model. As no fossils of Heliconiini or closely related tribes are known, we used a secondary calibration point from the dated phylogeny of Nymphalidae (Wahlberg et al. 2009; van Velzen et al. 2013). The age of the root was modelled as normally distributed with a mean of 47 Ma and a standard deviation of 3.0 Ma, corresponding to the 95% confidence intervals of the Acraeini-Heliconiini split found by Wahlberg et al. (2009). Four independent instances of the MCMC chain were ran for 100 million cycles each, sampling the posterior every 10000 cycles and discarding 10 million cycles as burnin. The input .xml file generated in BEAUti v. 1.7.5 (Drummond et al. 2010) can be found in the Online Appendix 3. To ensure that the results are driven by the data and not the priors, we executed a single empty prior run. Convergence of the continuous parameters was evaluated in Tracer v. 1.4, and the Maximum Clade Credibility tree with mean age of the nodes was generated using LogCombiner v. 1.7.5 and TreeAnalyser v. 1.7.5 (Rambaut and Drummond 2010).

### Multispecies Coalescent Phylogenetics

To account for the heterogeneous phylogenetic signal resulting from gene flow, hybridisation and incomplete lineage sorting, we applied a variety of multispecies coalescent (MSC) analyses that take as input both the raw alignment and individual gene trees). We first used the established method of minimising deep coalescences (MDC) (Maddison and Knowles 2006), taking advantage of a dynamic programming implementation in the package PhyloNet (Than et al. 2008). 100 bootstrap input files containing 100 trees were drawn randomly without replacement from the distribution of Bayesian gene trees for the 21 loci, an MDC phylogeny was estimated for each input and a 50% majority rule consensus was taken.

Bayesian Concordance Analysis (BCA) is an MSC method that attempts to reconcile the genealogies of individual loci based on posterior distributions, regardless of the sources of conflict (Larget et al. 2010). BCA generates Concordance Factors (CFs), which show what proportion of loci contain a particular clade, and estimates the primary phylogenetic hypothesis from the best-supported clades. CFs offer a powerful alternative to traditional measures of support and can be conveniently estimated in the program BUCKy (Ané et al. 2007; Larget et al. 2010). We executed two runs of one million MCMC cycles in BUCKy based on the 21 posterior distributions of gene trees from MrBayes.

Another approach to the multispecies coalescent is to estimate the gene trees and the species tree simultaneously, explicitly modelling the sources of incongruence (Edwards et al. 2007). We applied this technique using *BEAST, a program harnessing the power of BEAST and simultaneously implementing a powerful MSC algorithm that estimates the species tree and the embedded gene trees, as well as the population sizes of the lineages (Heled & Drummond 2010). Effective calculation of the population size parameters requires a thorough multilocus sampling of each species in the analysis, which forced us to reduce the dataset to species with a minimum of three individuals from at least two distinct populations (the <u>*BEAST dataset</u>). The final alignment included 87 terminal taxa in 17 species of *Eueides* and *Heliconius*. We re-estimated the individual substitution models for each partition, used a constant population size coalescent tree model and implemented other priors as described above for BEAST. We carried out four independent runs of 500 million cycles each, sampling every 10000 cycles, generated a maximum clade credibility species tree and visually summarised the 21 gene trees by plotting in DensiTree (Bouckaert 2010). Online Appendix 4 contains the .xml file for this analysis.

Sequence alignments and phylogenetic trees were deposited in TreeBase (http://purl.org/phylo/treebase/phylows/study/TB2:S15531).

### Changes in the Diversification Rate

The formal analyses of diversification dynamics were based on the output of the Bayesian supermatrix analysis in BEAST and conducted in R. Errors in divergence time estimates are not expected to affect the divergence rate calculations significantly (Wertheim and Sanderson 2011), and the semi-log lineage through time plots (LTT) of 200 randomly selected posterior trees formed a narrow distribution around the maximum clade credibility (MCC) tree with mean node ages. Hence we based all the analyses on the MCC tree of Heliconiini with one individual per species, excluding outgroups and *Cethosia*.

We searched for the signal of change in diversification rate using TurboMEDUSA, which identifies shifts *a posteriori* through stepwise AIC (Alfaro et al. 2009). We wanted to investigate whether the Heliconiini have undergone significantly faster diversification during the periods of ramatic environmental change at either 12 or 4.5 Ma, whether the diversification rate of Heliconiini decreased with time due to limiting species density effects (Fordyce 2010; Etienne et al. 2012b), or increased as new lineages evolved novel colour patterns and drove positive mimetic interactions that reduce competition (Elias et al. 2009). To compare these possibilities explicitly, we fitted maximum likelihood models of increasing complexity to the MCC tree, accounting for the uncertainty in the estimates of recent speciation (protracted speciation) and extinction rates (pull of the present) by truncating the phylogeny to 1 Ma before present (Etienne and Haegeman 2012). The following models were fitted and compared using the Akaike weights for all Heliconiini and for *Heliconius* alone: pure birth, with a constant rate or with a shift in rate (PB); birth-death with and without changes in rate (BD); diversity dependent model without extinction (DDL); diversity dependent model with linear terms for speciation or both speciation and extinction (DDE); DDE with rate shifts. We included rate shifts at an unspecified time, at 12 and 4.5 Ma, as well as at 11 and 3.5 Ma to account for possible delayed effects. The likelihood was conditioned on the survival of phylogeny in all runs and it was assumed that there are six species missing for Heliconiini and two for *Heliconius* (Lamas 2004). A possible slowdown in diversification rate after an initial rapid radiation was evaluated by calculating the gamma statistic (Pybus and Harvey 2000) and its significance tested in a Monte Carlo Constant Rates (MCCR) test with 1000 replicates in the package LASER (Rabosky 2006).

## Results

### Sampling and DNA Sequencing

We successfully combined three approaches to sequencing, which resulted in the high taxonomic coverage and intraspecific sampling necessary for the MSC methods, and a sufficient sampling of loci for each individual (Online Appendix 1). Most of the dataset consists of Sanger sequences from 108 individuals, 26 of which were already published in GenBank. We also obtained two classic lepidopteran markers *CoI/II* and *EF1α* from respectively 13 and 11 out of 14 historical specimens, thus adding eight species of Heliconiini that have not been sequenced previously. We did not manage to obtain any data from *Philaethria andrei*, *P. browni*, *P. romeroi*, *Eueides emsleyi*, *E. libitina*, *Heliconius lalitae* or *H. metis* (Lamas 2004; Constantino and Salazar 2010; Moreira and Mielke 2010).

We capitalised on the availability of Illumina data by generating *de novo* assembly contigs for further 57 individuals. The N50 of the Abyss assemblies ranged from 552 to 1921 bp (average 1206) and all the nuclear markers were successfully recovered by megaBLAST from every assembly. Whole mitochondrial sequences of the same individuals were recovered by read mapping, with about a 400 bp stretch of the hypervariable control region (Heliconius Genome Consortium 2012) incomplete in some sequences. We obtained a depth of coverage over 100x and high confidence in the base calls due to the high copy number of mtDNA in the tissue. Finally, we included the sequences extracted from the *Heliconius melpomene* genome, as well as the previously published mitochondrial sequence of *Acraea issoria* (Heliconius Genome Consortium 2012).

The sequence data for 20 nuclear and 2 mitochondrial genes encompass 71 out of 78 (91%) of the officially recognised species of Heliconiini, including 44 out of 46 species from the focal genus *Heliconius* (Lamas et al. 2004; Beltrán et al. 2007; Mallet et al. 2007; Constantino and Salazar 2010). Although the taxonomic validity of some species is contested, we found that the diversification analysis is robust to altering the number of missing species. Some recognised taxa diverged very recently, as shown by the BEAST chronogram (Fig. 1) in case of the *Philaethria diatonica/P. neildi/P. ostara* complex and the *Heliconius heurippa/H. tristero* pair, the latter of which could be considered races of *H. timareta* (Nadeau et al. 2013). However, the exact relationship between genetic differentiation and taxonomic species identity in the highly variable, mimetic Heliconiini remains unclear (e.g. Mérot et al. 2013). Importantly, 36 species are represented by multiple individuals, usually from distant populations, allowing for more accurate estimation of sequence evolution rates, and detection of species paraphyly. The number of individuals represented by each marker ranges from 40% for *Hcl* to 98% for *CoI/II*, and only four specimens are represented exclusively by mitochondrial DNA (Online Appendix 1).

**Figure 1.**
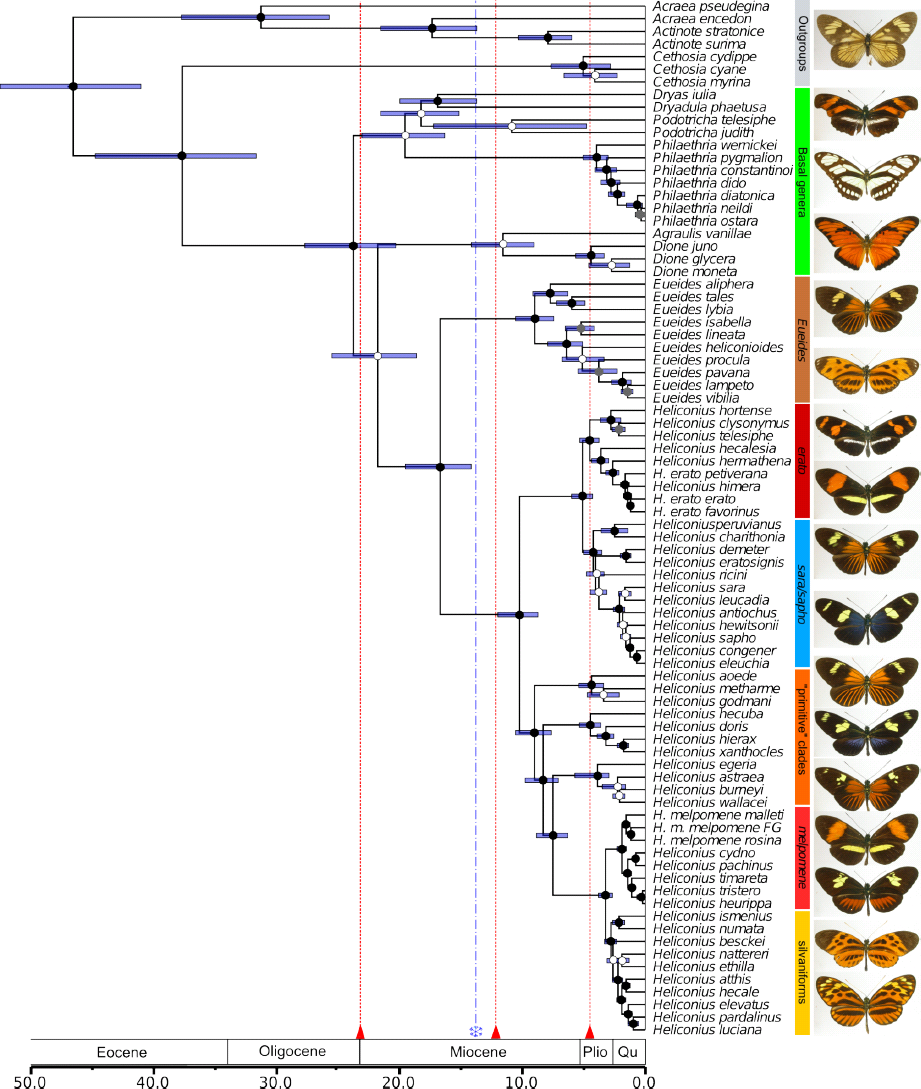
Bayesian phylogeny of 71 out of 78 butterflies in the tribe Heliconiini with outgroups, estimated using 22 markers with an uncorrelated molecular clock method (BEAST). The age of the root is calibrated based on the results of Wahlberg et al. (2009) and the bars signify the 95% confidence intervals around the mean node ages. The dots indicate the Bayesian posterior probability of the splits: >99% (black), 95-99% (grey), <95% (white). Scale axis in Ma. Red triangles indicate the periods of Andean orogenesis 23, 12 and 4.5 Ma and the blue asterisk marks the end of the Mid-Miocene Global Optimum (Hoorn et al. 2010). Deep splits are shown within the well-studied *Heliconius erato* and *H. melpomene*. FG=French Guiana. Heliconiini exhibit complex patterns of divergence and convergence in aposematic wing patterns, top to bottom: *Actinote latior* (outgroup), *Eueides tales michaeli*, *E. lampeto lampeto*, *H. telesiphe telesiphe*, *H. erato favorinus*, *H. demeter ucayalensis*, *H. sara*, *H. aoede*, *H. doris* (blue morph), *H. burneyi jamesi*, *H. melpomene amaryllis*, *H. timareta ssp.*, *H. numata lyrcaeus*, *H. pardalinus dilatus*.

### Pervasive Conflict Between the Loci

Our nuclear markers span 11 out of 21 chromosomes (Table 1; Heliconius Genome Consortium, 2012) and have both autosomal and sex chromosome Z-linked inheritance. We examined the conflict between individual markers in the entire tribe and in the genus *Heliconius* alone, using both gene tree summary methods and approaches utilising the raw sequence alignments. The maximum likelihood analysis of the <u>core matrix</u> in Concaterpillar rejected concatenation of any of the loci due to significant differences in both topology and substitution rate of individual partitions, but the exact nature of the discordance is unclear. A Multi-Dimensional Scaling ordination of pairwise RF distances between the gene trees does not reveal clustering by chromosome and the separation between many nuclear loci appears much greater than between nuclear and mitochondrial trees (Fig. 2). Consistent with this is the fact that the whole mitochondrial phylogeny of select taxa (Fig. 3) shows few differences from the tree based on the mixed marker supermatrix (Fig. 1), highlighting that cyto-nuclear discordance is not the primary source of incongruence in the dataset.

**Figure 2.**
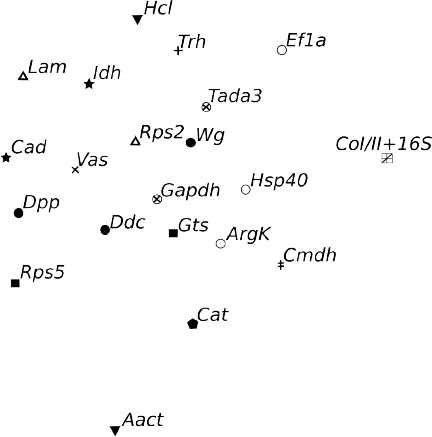
Multi-dimensional scaling ordination of pairwise Robinson-Foulds distances between 21 gene trees fitted in the Tree Set Visualiser shows no clear patterns of incongruence between the markers. Phylogenies of two mitochondrial and 20 nuclear markers were estimated in MrBayes. The axes have no units.

**Figure 3.**
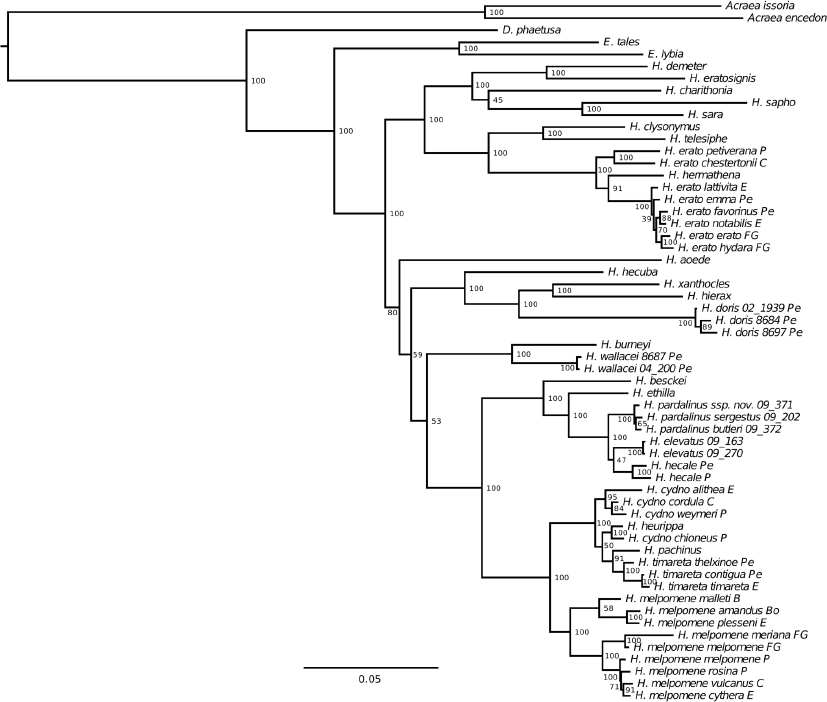
Whole mitochondrial maximum likelihood (RAxML) phylogeny of the genus *Heliconius*. Bootstrap support values indicated. Scale bar in units of substitution per site per million years. B=Brazil, Bo=Bolivia, C=Colombia, E=Ecuador, FG=French Guiana, P=Panama, Pe=Peru.

The coalescent approaches reveal the high extent of marker conflict in the *Heliconius* data. Gene tree topologies from the explicit Bayesian modelling of incomplete lineage sorting (ILS) in *BEAST are highly varied, with a particularly high degree of reticulation in the *H. melpomene/cydno* and the *H. hecale* (silvaniform) clades, where extensive horizontal gene flow has been observed previously (Sup. Fig. S1; Brown 1981; Martin et al. 2013). Another Bayesian method, BUCKy, infers the species in the presence of marker incongruence without modelling specific reasons for the observed discordance and calculates the Bayesian concordance factors that illustrate the proportion of partitions in the dataset that support a particular grouping (Ané et al. 2007; Baum 2007; Larget et al. 2010). The concordance of the loci for *Heliconius* is strikingly low (Fig. 4), although the topology is consistent with the results of other analyses (Fig. 1 and 3, Sup. Fig. S2-S6).

**Figure 4.**
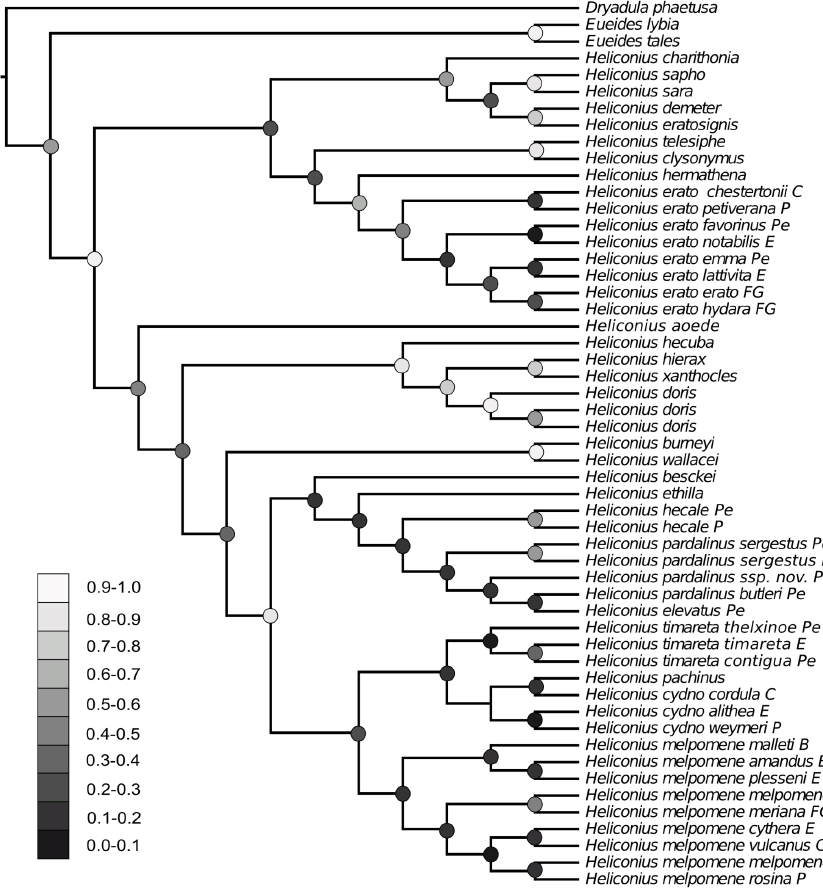
A phylogenetic hypothesis for *Heliconius* showing the extent of concordance between tree topologies of the 21 loci estimated by Bayesian concordance analysis in BUCKy. Dots indicate the Concordance Factor values for the nodes, with darker shades of grey corresponding to lower support values. B=Brazil, Bo=Bolivia, C=Colombia, E=Ecuador, FG=French Guiana, P=Panama, Pe=Peru.

Further strong evidence of widespread incongruence comes from the NeighborNet network characterised by a high delta score of 0.276, which shows that the structure of the data is not entirely tree-like (Sup. Fig. S3). This can be partially attributed to the effect of missing data, yet even a fit based on the 30 species with little missing data produced a delta score of 0.11, proving a substantial amount of non-bifurcating signal across the tribe (Holland et al. 2002). The most noticeable reticulations are found between nodes linking genera and the major clades of *Heliconius*, possibly due to pervasive gene flow during the diversification of the main extant lineages (Sup. Fig. S3).

### Topological Consistency Across Optimality Criteria

Although no trees are identical, the results from our Bayesian, Maximum Likelihood and distance-based network analyses of the supermatrix are very similar (Fig. 1, Sup. Fig. S3, S4). Two nodes stand out as unstable. *Cethosia* is variably placed as a sister taxon to either Acraeini (MP, ML, NeighborNet) or Heliconiini (Bayesian), despite the reasonably extensive sampling of 11 loci for *C. cyane*, while the position of *Podotricha* in relation to *Dryas* and *Dryadula* varies between all analyses. Most problematic are relations among the species within *Eueides*, where the position of four out of 10 taxa cannot be resolved with good support. The poor resolution for both *Eueides* and *Podotricha* can probably be attributed to insufficient site coverage, which produces high uncertainty due to patchily distributed missing data (Wiens and Morrill 2011; Roure et al. 2013). We reestimated the relations of *Eueides* based solely on a core set of 11 genes with coverage for at least 7/10 species and recovered a much better supported tree (Sup. Fig. S6). We recommend studies focusing on *Eueides* use this specific phylogeny, but the exact relations of *E. procula*, *E. lineata* and *E. heliconioides* remain unclear.

Although our Bayesian maximum clade credibility tree is largely consistent with the topology estimated by Beltrán *et al.* (2007), significantly increasing the dataset from 113 to 180 individuals and five to 21 loci allowed us to resolve many critical nodes. The major differences are in the relations of the basal genera, which we infer to form a well-supported grade, with *Eueides* as the definitive sister genus of *Heliconius* (Fig. 1). Importantly, we confirm that the enigmatic genera *Laparus* and *Neruda* are nested within *Heliconius*, as further supported by ELW and SH tests (Table 2), although *Neruda* is closer to the base of the genus than previously estimated. We find that the other so called “primitive” (Brown 1981) clade of *Heliconius* consists of two separate groups, which are by no means basal to the other taxa, raising questions about the apparently unequal rate of morphological evolution in the genus.

**Table 2.** Maximum likelihood tests of alternative topologies. Phylogenies were estimated under constraints enforcing alternative topologies and compared to the original ML tree with the Shimodaira-Hasegawa test and the Expected Likelihood Weights test with 100 bootstrap replicates.

| Grouping | Likelihood | ELW probability | SH result |
| --- | --- | --- | --- |
| ML tree | -142583 | 0.117 |  |
| *Cethosia*+*Heliconiini* | -142594 | 0.048 |  |
| *Agraulis*+*Dione* basal | -142599 | 0.006 |  |
| basal genera monophyletic | -142596 | 0.025 |  |
| *Dryas*+*Dryadula* monophyletic | -142593 | 0.058 |  |
| *Neruda* not in *Heliconius* | -142636 | 0.000 | *p*<0.05 |
| *Laparus* not in *Heliconius* | -143233 | 0.000 | *p*<0.05 |
| *H. erato* monophyletic | -142592 | 0.114 |  |
| *H. ricini* basal to *H. demeter* | -142590 | 0.084 |  |
| *H. antiochus* basal to *H sara* | -142588 | 0.170 |  |
| primitive clade monophyletic | -142613 | 0.000 | *p*<0.05 |
| *H. wallacei* monophyletic | -142584 | 0.115 |  |
| *H. elevatus* monophyletic | -142590 | 0.041 |  |
| *H. cydno* monophyletic | -142619 | 0.001 | *p*<0.05 |
| *H. heurippa* with *H. timareta* | -142586 | 0.209 |  |
| *H. melpomene* East+Guiana clade | -142620 | 0.011 | *p*<0.05 |

The maximum likelihood tree is similar to the Bayesian phylogeny, although characterised by lower overall support, especially for the nodes linking the genera and subgenera (Sup. Fig. S4). The nodes differing between the two trees are also the nodes that cannot be unequivocally confirmed by either the SH and the ELW test (Table 2). For instance, the highest posterior probability in the ELW test (0.21) is given to a tree placing *H. heurippa* as a sister species to *H. timareta*, a result recovered in both the Bayesian analysis and a recent genome-wide study (Nadeau et al. 2013). Thus we suggest that the Bayesian tree should be preferred as a more accurate picture of the phylogenetic relationships, although the ML tree based on the <u>complete</u> dataset is still useful to uncover multiple polyphyletic species. Notably, *H. luciana* is nested within *H. elevatus*, and *H. wallacei i*s polyphyletic with respect to *H. astraea*. These results must be interpreted with caution, as the inference relies on poorly covered museum specimens and may be sensitive to long branch attraction (Wiens and Morrill 2011).

Our whole mitochondrial phylogeny is largely consistent with the results of the multilocus supermatrix analysis and well-supported for 46/57 nodes (Fig. 3), despite the relatively limited taxonomic coverage of only 29 Heliconiini for which short-read data was available. In contrast to the multilocus dataset, mitochondrial genomes are not very useful for resolving relationships between major clades within *Heliconius*, as the position of *Neruda*, *xanthocles* and *wallacei* clades is poorly supported. An important deviation from the predominantly nuclear multilocus phylogeny is the placement of *H. pachinus* as a sister taxon of *H. timareta*, rather than *H. cydno*. This may reflect the overall instability resulting from a high extent of reticulation in the *melpomene/cydno/timareta* assemblage (Heliconius Genome Consortium 2012, Martin et al. 2013, Mérot et al. 2013). Furthermore, we find a surprising positioning of *H. hermathena* within *H. erato* (Jiggins et al. 2008), which is also not supported by any of the analyses of the 21 locus matrix. The mitochondrial tree confirms previous observations of deep biogeographical splits in the widespread, highly diversified *H. erato* and its co-mimic *H. melpomene* (Brower 1994a). Within *H. melpomene* we find a well-supported distinction between races found to the West and East of the Andes, although our data place the individuals from French Guiana with the specimens from the Western clade, in contrast to a whole-genome phylogeny (Nadeau et al. 2013) (Fig. 1). *H. erato* shows the opposite pattern in both mitochondrial and nuclear data, whereby the Guianian races form a fully supported clade in the monophyletic group of taxa from East of the Andes.

The variety of multi-species coalescent (MSC) methods that we applied brings a new perspective to the phylogenetic signals in the Heliconiini. Notably, different approaches and combinations of the data yield highly consistent topologies, although the support for the individual nodes varies. We used three techniques that attempt to deal with various aspects of the observed incongruence between loci. The maximum parsimony approach of MDC is a summary method deriving a species tree from the distribution of gene trees with single or multiple terminals per species, intended to address the effects of ILS (Maddison 1997; Maddison and Knowles 2006). Although infrequently considered in recent studies, MDC has been implemented in multiple packages (Maddison and Maddison 2011; Than et al. 2008) and offers superior speed of analysis when compared to other MSC techniques. In case of our <u>complete</u> dataset, a run with 100 bootstrap datasets took a few minutes. The resulting topology is nonetheless very different from the other results, showing a number of unexpected and poorly supported groupings. The lumping of all non-*Heliconius* genera, and the monophyletic *Neruda/xanthocles/wallacei* clade stand out in contrast to other proposed trees (Sup. Fig. S6). Interestingly, many of the relations that are poorly supported in the supermatrix phylogenies are also not resolved in the consensus MDC tree, showing that MDC is highly conservative with regard to the placement of taxa unstable in individual gene trees.

The Bayesian MSC approach implemented in *BEAST uses MCMC to co-estimate the species-level tree and the contained gene trees by estimating demographic parameters together with the phylogeny, but only with an appropriate intraspecific sampling (Heled and Drummond 2010). Although computationally intensive, *BEAST models ILS and proposes a species tree without ignoring the underlying heterogeneity in specific locus genealogies, performing exceptionally well even with the most difficult cases combining large population size with short divergence times (Leaché & Rannala 2010). We analysed the small dataset of the 17 best-sampled species and recovered a tree which agrees with the supermatrix analyses (Sup. Fig. S2), except for the position of *Neruda aoede*, which was placed as a sister taxon to the *H. xanthocles/L. doris* clade with relatively low posterior probability. Furthermore, the mean ages of nodes are nearly identical to those proposed in a supermatrix analysis, with similar 95% confidence intervals. Although the species tree is largely as predicted, we observe high levels of incongruence in the underlying distribution of gene trees (Sup. Fig. S1). Differences in the depth of coalescence are clear throughout the tree and reticulation is again especially apparent in the *H. melpomene/cydno* clade. The estimated population size values are also consistent with a previous comparison based on two nuclear loci, showing a higher population size of *H. erato* (1.33×10^6^ individuals) when compared to *H. melpomene* (1.02×10^6^) (Flanagan et al. 2004).

While *BEAST is a powerful approach to account for ILS under a complex evolutionary model, it does not take into account various other sources of data heterogeneity. BUCKy derives a topology together with CFs, which represent the proportion of the genome supporting a particular clade. The most likely clade may therefore receive low support if it is represented in only a few of the Bayesian gene tree posteriors (Larget et al. 2010). The method is thus able to propose a robust topology in the face of ILS, hybridisation or other complex processes. As described above, we find the phylogeny derived by BUCKY to be entirely consistent with the Bayesian analysis of concatenated sequence, although the recovered CFs are much lower than any other measure of support applied to our data. Importantly, most of the nodes connecting the major clades in the tree have CFs below 0.5, with the notable exception of the silvaniform/*melpomene* split (Fig. 4). The same nodes correspond to the reticulations in the NeighborNet analysis (Sup. Fig. S3), cases of low support in the MDC tree and its disagreement with the supermatrix analysis (Sup. Fig. S6), nodes that cannot be rejected in the ML tests of topologies (Table 2), and the uncertain nodes in the whole mitochondrial tree (Fig. 3).

### Tempo of Diversification

The phylogeny estimated under a relaxed clock model in BEAST shows diversification dynamics that differ from previous estimates, with the splits between the genera of Heliconiini being older and the extant species and subgenera much younger than previously thought (Table 3)(Mallet et al. 2007). The most speciose genera *Heliconius* and *Eueides* separated 17 Ma and both started to diversify around 10 Ma, and the six major clades of *Heliconius* (corresponding to *H. erato*, *H. sara*, *H. xanthocles*, *H. wallacei*, *H. melpomene*/silvaniforms and *Neruda*) all started to diversify between 5 and 4 Ma (Fig. 1). The lineage through time plot (LTT) for Heliconiini suggests a period of stasis corresponding to the mid-Miocene 16-11 Ma (Hoorn et al. 2010), and followed by a sudden increase in the number of extant lineages (Fig. 5a). In case of the 45 *Heliconius* species, a shorter slowdown is found between 7 and 4.5 Ma (Fig. 5b) (Hoorn et al. 2010). As expected from the LTT plots, the MCCR test detects no evidence for an overall slowdown in the diversification rate (Heliconiini: gamma= 2.643, p=0.996; *Heliconius*: gamma= 0.153, p=0.617).

**Table 3.** Mean split ages in Ma at different levels of divergence within Heliconiini, as estimated by previous studies and by two Bayesian relaxed clock methods in the present work. Mean ages and 95% confidence intervals are reported where available.

|  | *Heliconius*<br>vs *Eueides* | *Heliconius* vs<br>*Agraulis* | *Heliconius* vs<br>*Philaethria* | *H. melpomene*<br>vs *H. erato* | *H. erato* vs<br>*H. hecalesia* |
| --- | --- | --- | --- | --- | --- |
| Mallet et al.<br>2007 | 11.0 | 13.0 | 14.5 | 9.5 | 4.0 |
| Pohl et al.<br>2009 | n/a | 32.0<br>(40.0-24.0) | n/a | 17.0<br>(11.0-23.0) | n/a |
| Wahlberg et<br>al. 2009 | 18.5<br>(12.5-24.5) | 26.5<br>(21-32) | 30.0<br>(36.5-23.5) | n/a | n/a |
| Cuthill and<br>Charleston<br>2012 | 13.0<br>(10.5-16.5) | n/a | n/a | 10.5<br>(8-13.5) | 4.5<br>(2.8-6.3) |
| BEAST | 16.7<br>(14.1-19.5) | 21.8<br>(18.6-25.5) | 23.8<br>(20.3-27.7) | 10.2<br>(8.7-12.0) | 3.6<br>(3.0-4.4) |
| *BEAST | 17.5<br>(11.6-23.4) | n/a | n/a | 10.6<br>(14.9-6.7) | n/a |

**Figure 5.**
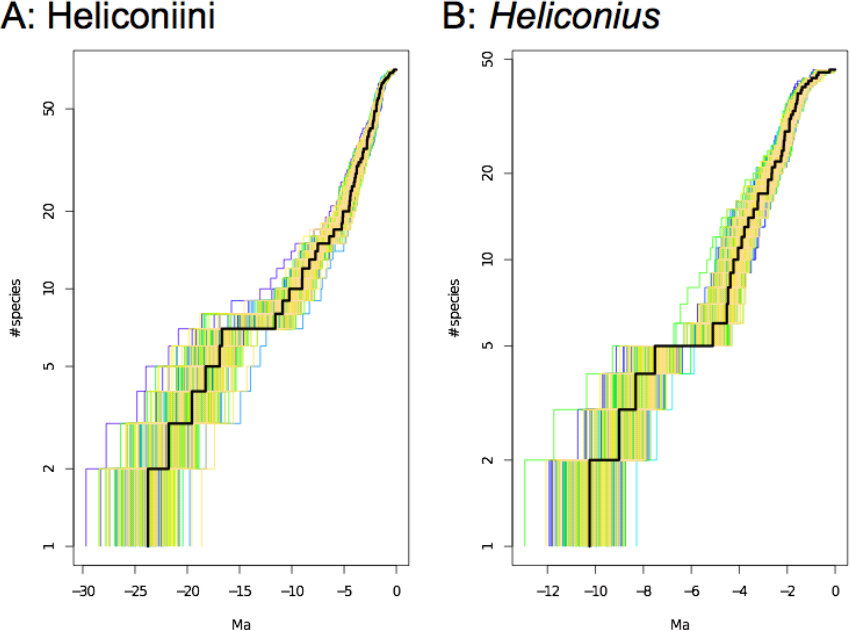
Lineage through time (LTT) plots based on the calibrated, uncorrelated molecular clock analysis in BEAST. A. Semilogarithmic LTT plot of the Heliconiini butterflies with the maximum clade credibility phylogeny in black and a random sample of 200 posterior trees around. B. LTT plot for *Heliconius*.

Maximum Likelihood (ML) modelling strongly supports the Birth-Death model with rate shift as the best fit for Heliconiini (Akaike weight of 0.92; Table 4). Both DDD and turboMEDUSA models demonstrate that around 11 Ma the speciation rate of Heliconiini increased dramatically from 0.19 to 0.40 new species per lineage per million years, and the shift was accompanied by an increase in the turnover rate from 0.11 to 0.65. The results for *Heliconius* are less clear, as equal Akaike weights (0.30) are given to the Pure Birth models without rate changes, and with an almost twofold increase in speciation rate 4.5 Ma (Table 4). TurboMEDUSA and DDD models allowing for a shift at an unspecified time also find that speciation accelerated 10-11 Ma and 4-5 Ma.

**Table 4.** Results of diversification model fitting to the maximum clade credibility chronograms of Heliconiini and *Heliconius*, estimated under a relaxed molecular clock model in BEAST. Models are compared in terms of corrected AIC (AICc) and relative Akaike weights (wAIC). Models: PB=Pure Birth, BD=Birth-Death, DD-E=Diversity Dependence without Extinction, DD+E1=Diversity Dependence with Extinction and a linear scaling of speciation term, DD+E5=Linear Dependence With Extinction and linear scaling terms for speciation and extinction. Where indicated, models include shifts in speciation and extinction rates at fixed times, or at any time.

| Model | $\lambda$ | $\lambda_2$ | $\mu$ | $\mu_2$ | K | K<sub>2</sub> | -lnL | AICc | wAIC |
| --- | --- | --- | --- | --- | --- | --- | --- | --- | --- |
| **Heliconiini** |  |  |  |  |  |  |  |  |  |
| PB | 0.23 |  |  |  |  |  | -158.48 | 319.01 | 0.00 |
| PB, shift 3.5 Ma | 0.17 | 0.31 |  |  |  |  | -155.43 | 315.04 | 0.00 |
| PB, shift 4.5 Ma | 0.13 | 0.33 |  |  |  |  | -152.27 | 308.72 | 0.03 |
| PB, shift 11 Ma | 0.11 | 0.26 |  |  |  |  | -155.57 | 315.32 | 0.00 |
| PB, shift 12 Ma | 0.11 | 0.25 |  |  |  |  | -155.82 | 315.81 | 0.00 |
| PB, any shift | 0.12 | 0.32 |  |  |  |  | -151.25 | 308.86 | 0.03 |
| BD | 0.38 |  | 0.26 |  |  |  | -154.62 | 313.41 | 0.00 |
| BD, shift 3.5 Ma | 0.31 | 0.31 | 0.23 | 0.00 |  |  | -153.50 | 315.61 | 0.00 |
| BD, shift 4.5 Ma | 0.16 | 0.33 | 0.04 | 0.00 |  |  | -152.27 | 313.14 | 0.00 |
| **BD, shift 11 Ma** | **0.19** | **0.40** | **0.02** | **0.26** |  |  | **-146.70** | **302.00** | **0.92** |
| BD, shift 12 Ma | 0.21 | 0.37 | 0.02 | 0.25 |  |  | -151.85 | 312.32 | 0.01 |
| BD, any shift | 0.26 | 0.37 | 0.16 | 0.23 |  |  | -153.82 | 318.56 | 0.00 |
| DD-E | 0.23 |  |  |  | Inf |  | -158.48 | 321.13 | 0.00 |
| DDE1 | 0.35 |  | 0.20 |  | 16x10<sup>5</sup> |  | -154.27 | 314.90 | 0.00 |
| DDE5 | 0.36 |  | 0.24 |  | 2190 |  | -153.64 | 313.63 | 0.00 |
| DDE1, shift 3.5 Ma | 0.31 | 0.42 | 0.23 | 0.00 | 7912 | 208 | -153.41 | 320.13 | 0.00 |
| DDE1, shift 4.5 Ma | 0.15 | 0.45 | 0.04 | 0.01 | Inf | 171 | -151.97 | 317.25 | 0.00 |
| DDE1, shift 11 Ma | 0.94 | 0.75 | 0.40 | 0.38 | 33 | 68 | -155.09 | 323.49 | 0.00 |
| DDE1, shift 12 Ma | 0.17 | 0.30 | 0.00 | 0.07 | 26 | 941 | -154.36 | 322.02 | 0.00 |
| DDE1, any shift | 0.16 | 0.49 | 0.01 | 0.07 | 71 | 152 | -151.18 | 318.13 | 0.00 |
| *Heliconius* |  |  |  |  |  |  |  |  |  |
| **PB** | **0.42** |  |  |  |  |  | **-73.87** | **149.82** | **0.30** |
| PB, shift 3.5 Ma | 0.38 | 0.44 |  |  |  |  | -73.77 | 151.82 | 0.11 |
| **PB, shift 4.5 Ma** | **0.26** | **0.47** |  |  |  |  | **-72.79** | **149.86** | **0.30** |
| BD | 0.43 |  | 0.02 |  |  |  | -73.86 | 152.00 | 0.10 |
| BD, shift 3.5 Ma | 0.60 | 0.44 | 0.39 | 0.00 |  |  | -73.26 | 155.50 | 0.02 |
| BD, shift 4.5 Ma | 0.26 | 0.47 | 0.00 | 0.00 |  |  | -72.79 | 154.56 | 0.03 |
| DD-E | 0.42 |  |  |  | Inf |  | -73.87 | 152.01 | 0.10 |
| DDE1 | 0.45 |  | 0.07 |  | 6x10<sup>5</sup> |  | -73.88 | 154.34 | 0.03 |
| DDE5 | 0.43 |  | 0.02 |  | Inf |  | -73.86 | 156.70 | 0.01 |
| DDE1, shift 3.5 Ma | 0.39 | 0.60 | 0.02 | 0.00 | 11x10<sup>6</sup> | 111 | -73.51 | 161.16 | 0.00 |
| DDE1, shift 4.5 Ma | 0.26 | 0.70 | 0.00 | 0.00 | Inf | 85 | -72.18 | 158.51 | 0.00 |
| DDE1, any shift | 0.38 | 0.45 | 0.01 | 0.00 | 128 | 9710 | -73.79 | 164.53 | 0.00 |

## Discussion

### Stable Topology Despite Marker Incongruence

We approached the problem of phylogeny reconstruction in a difficult mimetic assemblage through extensive intraspecific sampling of 22 markers from nearly all species in the clade, and compared the results between multiple philosophically distinct analytical approaches. As next generation sequencing technologies become widely accessible and the average number of loci used in systematic studies increases rapidly, Multi-Species Coalescent (MSC) methods gain in importance as means of detecting and accounting for incongruence in multilocus data (e.g. Lee et al. 2012; Barker et al. 2013; Smith et al. 2013). However, their relative merits and utility at different levels remains contested (Song et al. 2012; Gatesy and Springer 2013; Reid et al. 2013). The systematic relations of the tribe Heliconiini, which diverged from its extant outgroup about 47 Ma, can be effectively resolved with both MSC and supermatrix approaches, yielding highly similar topologies across a range of different sub-sampling schemes that correspond to the requirements of individual algorithms (Fig. 1, 3 and 4; Sup. Fig. S2-S6). Nonetheless, consistent with the recent radiation of the group, large effective population sizes and known hybridisation between many species, we observed high heterogeneity among sampled fragments of the genome that differ markedly in both topology (Fig. 2, Sup. Fig. S1) and rates of evolution (Concaterpillar analysis). Such heterogeneity might have been expected to pose a significant challenge for the concatenation methods (Degnan and Rosenberg 2006; Edwards et al. 2007; Edwards 2009; Knowles and Kubatko 2010; Leaché and Rannala 2010). However, the only method producing an obviously different phylogeny is Minimising Deep Coalescences (MDC; Sup. Fig. S6), which fails to resolve 34 out of 62 nodes in the tree with bootstrap support above 0.9, and is the only method to suggest monophyly of *Eueides* with other basal genera of Heliconiini. MDC derives a species tree from point estimates of gene trees and can be expected to perform poorly with a relatively limited number of gene trees that are not always fully resolved, leading to a complete polytomy in some of the clades (Gatesy and Springer 2013). However, the MDC result is an indicator of instability, as well-resolved and consistent gene trees should produce a good quality MDC tree.

Recent studies have proposed that likelihood-based MSC techniques perform better than and should be preferred to integration over individual gene trees, due to their potential to capture synergistic effects between partitions (Leaché and Rannala 2010; Reid et al. 2013), mirroring the phenomenon of hidden support found when multiple markers are concatenated (Gatesy and Baker 2005). Our results support this observation and further show that high degrees of conflict between many partitions can be reconciled by both supermatrix and MSC approaches to extract the predominant signal of speciation. The superiority of the MSC methods lies in the effective demonstration of incongruences, represented by lower support values or concordance factors assigned to the more difficult nodes (Belfiore et al. 2008). The Bayesian concordance analysis in BUCKy assigns insignificant concordance factors to most of the nodes separating major subgenera of *Heliconius*. Two of these nodes (*H. wallacei* and *N. aoede*; Fig. 4) are also only weakly supported by the *BEAST coalescent model, and correspond to the areas of high reticulation in the NeighborNet network, reflecting conflicting signals (Sup. Fig. S3). The same nodes are all assigned a posterior probability of one in the Bayesian supermatrix analysis (Fig. 1), potentially leading to the erroneous conclusion that all the data point unequivocally to the inferred relations.

Our study highlights an important practical consideration in choosing the optimal analytical approach, where the requirements of the selected algorithm have to be reconciled with a realistic sampling of taxa. Our final goal of testing hypotheses regarding the diversification dynamics of Heliconiini can be met only if the number of included taxa is maximised. Despite a substantial effort we only managed to secure single samples of the rare or geographically restricted species, and some of them are represented solely by historical specimens with limited potential to generate extensive multilocus data. Considering that our study group is intensively studied and exceptionally well represented in research, museum and private collections due to its aesthetic appeal (Mallet et al. 2007), it would be considerably more challenging to obtain a complete sampling of many other groups. Another difficulty stems from the fact that the advanced coalescent techniques like BUCKy and *BEAST perform best with multiple samples per species, which should capture intraspecific diversity (Heled & Drummond 2010, Reid et al. 2013). Finally, much of the uncertainty in the estimates can be attributed to missing data, which can negatively affect the estimation of both individual gene trees and the encompassing species tree (Wiens and Morrill 2011; Roure et al. 2013). When fitting a NeighborNet network, we found that although the percentage of data missing from the matrix does not explain all of the observed reticulation and the high delta score, it causes these parameters to increase, thus suggesting that the data completeness at each alignment position must be maximised to identify genuine incongruence. We observe that in many cases of biological interest it will be a formidable challenge to generate the ideal dataset that (i) has little missing data, (ii) comprises a large, genome-wide sample of loci, (iii) includes all taxa, and (iv) captures intraspecific variability. In case of Heliconiini, the supermatrix approach based on a limited number of markers (22) helped us to maximise taxonomic inclusiveness without compromising our ability to reconstruct a phylogeny in the light of conflicting biological signals.

### Divergence time estimates

Our phylogeny of Heliconiini brings novel insight into the diversification dynamics of the clade. Although most age estimates for *Heliconius* agree with other studies, the deeper nodes are older than previously suggested (Table 3). There is little agreement on the dates above the species level, and the studies to date either suffer from insufficient taxon sampling (Pohl et al. 2009, Wahlberg et al. 2009, Cuthill and Charleston 2012; Table 3), or use markers unlikely to be informative above a relatively low level of divergence (Mallet et al. 2007). For instance, the mean age of the split between *Heliconius* and *Agraulis* is estimated as 32 Ma in the study of Pohl et al. (2009), which includes only 3 species of Heliconiini; 26.5 Ma in Wahlberg et al. (2009), including one species per genus; or 21 Ma in the present study (Fig. 1; Table 3). Conversely, Mallet et al. (2007) find the divergences between the basal genera of Heliconiini to be much younger than we propose, likely due to an effect of using a fast-evolving mitochondrial locus without partitioning (Brandley et al. 2011).

Our own ability to correctly falsify the hypotheses regarding the diversification of Heliconiini hinges on having a nearly complete phylogeny, yet none of our MSC analyses consider as many species of Heliconiini as the Bayesian supermatrix estimate. We are confident that the values proposed by the supermatrix method can be trusted, as both the topology and the branch lengths are consistent with the results inferred by *BEAST based on a smaller dataset (Table 3; Sup. Fig. S2). We find that the ages of the deeper nodes and the length of terminal branches are not inflated by the supermatrix method in comparison to *BEAST, contrary to the predictions from simulations (Burbrink and Pyron 2011), and both methods infer similar mean age for the observed splits, for instance 10.5 Ma for the basal divergence of *Heliconius* into the *H. erato* and *H. melpomene* lineages, or 2-3 Ma for the diversification of *H. melpomene* and silvaniform clades. Hence we offer a new perspective on the dating of Heliconiini radiation with a nearly complete set of divergence time estimates based on a well-resolved and supported tree.

### Rapid Adaptive Radiation of *Heliconius*

*Heliconius* have undergone a rapid adaptive radiation *sensu* Schluter (2000), associated with the evolution of Müllerian mimicry, close coevolution with the Passifloraceae host plants (Benson et al. 1975) and other traits unique among the Lepidoptera (reviewed by Beltrán et al. 2007). Maximum Likelihood models demonstrate that around 11 Ma the speciation rate of Heliconiini increased dramatically from 0.19 to 0.40 new species per lineage per million years, and the shift was accompanied by an increase in the turnover rate from 0.11 to 0.65, indicating higher likelihood of extinction of the lineages. Our results are consistent with the radiation having been stimulated by environmental factors such as mountain uplift, and creation of elevation gradients facilitating parapatric speciation (Wesselingh et al. 2009, Hoorn et al. 2010, Turchetto-Zolet et al. 2013). Consistent with the inference of the Central and Eastern Andean slopes as the “species pump” of Heliconiini (Rosser et al. 2012), we find that diversification of *Heliconius* and *Eueides* into subgeneric clades occurred within the two million years after the major Andean uplift event 12 Ma, and that the modern species of *Heliconius* started to diversify after the last intensive orogenic event 4.5 Ma (Gregory-Wodzicki 2000). Contrary to our expectations, mimicry does not leave a clear signature of positive feedback on the rate of diversification. Although mimetic interactions have been repeatedly invoked as drivers of speciation in Heliconiini and butterflies generally (Bates 1862; McMillan et al. 1997; Mallet and Joron 1999; Elias et al. 2008; Jiggins 2008; Savage and Mullen 2009) and mimicry can possibly decrease the risk of extinction as a consequence of reduced predation rates (Vamosi 2005), model fitting does not support an exponential increase in rates of speciation.

Although we find sudden increases in speciation rates, there is no indication that the rates slowed down significantly afterwards. This result is consistent with a recent critique (Day et al. 2013) of the traditional prediction that large radiations should display a pattern of an initially high net diversification rate, decreasing as the ecological niches fill up (e.g. Rabosky and Lovette 2008). This expectation is reasonable for spatially limited island radiations that constitute the majority of study cases to-date, but it does not necessarily apply to continental radiations in the tropics, where the scale and complexity of the ecosystems are likely to generate a number of suitable niches greatly exceeding even the cladogenetic potential of large radiations (Day et al. 2013). So far, steady diversification of a widely distributed taxon has been demonstrated in the African *Synodontis* catfish (Day et al. 2013) and the Neotropical Furnariidae ovenbirds (Derryberry et al. 2011), but the generality of this pattern remains unknown. Heliconiini constitute an interesting case of a widespread, continental radiation where at least one sharp transition to a higher species turnover punctuates an otherwise steady diversification. Thus the physical environment has acted a driver of diversification, but did not limit cladogenesis.

Some uncertainty surrounds the last few million years of evolution of Heliconiini. Early speculation regarding the drivers of speciation suggested diversification in allopatry, as the rainforest habitat occupied by most species in the group has undergone cycles of contraction and expansion in response to recent climatic variation (Turner 1965; Brown et al. 1974; Brown 1981; Sheppard et al. 1985; Brower 1994a). The hypothesis of vicariant cladogenesis has been subsequently criticised due to a lack of evidence for forest fragmentation in pollen core data (Colinvaux et al. 2000, Dasmahapatra et al. 2010), as well as the likelihood of parapatric speciation (Mallet et al. 1998). The decrease in observed diversification over the last million years is likely due to our limited ability to delineate species in the assemblages of highly variable taxa like *Heliconius* – a phenomenon that has been termed “protracted speciation” (Etienne & Haegemann 2012). We provide direct evidence that most specific divergences between species of Heliconiini occurred during the Miocene, and that both climate and orogeny strongly influenced the pattern of diversification, although we cannot directly test the causes of this association with these data.

### Taxonomic and Systematic Implications

Application of novel algorithms to the Heliconiini data reveals a relatively stable topology despite limited support. We uphold the previous re-classification of the species in the genera *Neruda* (Huebner 1813) and *Laparus* (Linnaeus 1771) as *Heliconius* based on their nested position (Beltrán et al. 2007). This placement is at odds with morphological evidence from adult and larval characters, which puts *Neruda* and *Laparus* together with *Eueides* (Penz 1999), suggesting convergent evolution of homoplasious morphological characters. Nevertheless, the molecular evidence is decisive and we thus synonymize *Laparus* **syn. nov.** and *Neruda* **syn. nov.** with *Heliconius*.

The position of the enigmatic genus *Cethosia* remains unresolved, as it currently depends on the chosen method of analysis, and is unlikely to be established without a broad sampling of species using multiple markers. *Cethosia* has been variably considered to be either the only Old World representative of Heliconiini (Brown 1981; Penz and Peggie 2003; Beltrán et al. 2007), a genus of Acraeini (Penz 1999; Wahlberg et al. 2009), or possibly a distinct tribe (Müller and Beheregaray 2010). Establishing the systematic relations between Acraeini, *Cethosia* and Heliconiini is important for the study of Heliconiini macroevolution, as it could shed light on the origins of the heliconian affinity for the Passifloraceae host plants. Known records indicate that *Cethosia* species feed on *Adenia* and *Passiflora* (Müller and Behergaray 2010), raising the possibility that Heliconiini may have evolved from an Australasian ancestor that already exhibited the modern dietary preference.

Heliconiini have been at the centre of the debate about species concepts and designation criteria, providing empirical evidence for the permeability of species barriers (Mallet et al. 2007, Nadeau et al. 2012, Martin et al. 2013). At the same time, new species continue to be described based on morphological, genetic or karyotypic evidence (Lamas et al. 2004; Constantino and Salazar 2010; Dasmahapatra et al. 2010; Moreira and Mielke 2010), although the validity of the taxonomic status of some of these new lineages is disputed (J. Mallet and A. Brower, *pers. comm.*). Our diversification analysis is therefore prone to error resulting from the uncertainty in the number of terminal taxa that should be considered. We assume the number of species in the clade to be 78, reflecting the published taxonomy (Lamas 2004; Constantino and Salazar 2010; Rosser et al. 2012), and we deliberately exclude a formally undescribed species of *Agraulis*. Although we included *H. tristero* at the time of our analysis, it is not considered a species any longer (Mérot et al. 2013). In addition, a recent study suggests that Philaethria *pygmalion* and *Philaethria wernickei* may in fact constitute a single species (Barão et al. 2014). Although the number of Heliconiini species may therefore vary between 73 and 79, changing the number of missing species in the diversification analyses does not alter the results substantially.

### Conclusion

We present a taxonomically comprehensive phylogeny of a large continental radiation characterised by extensive hybridisation, adaptive introgression and rapid speciation, and emerging as a prominent system for comparative genomics. The tribe Heliconiini has radiated in association with environmental change in the Neotropics over the past 23 Ma and that speciation and species turnover increased dramatically at the end of the Mid-Miocene. Although there is a clear signature of processes leading to incongruence of individual gene trees and the species tree, we consistently recover the same topology using different analytical approaches. We conclude that the established supermatrix methods perform well, but are less effective in detecting the underlying conflict and estimating nodal support than the alternative Multispecies Coalescent techniques.

Müllerian mimicry affects the mode of speciation in *Heliconius*, and probably plays a role for the other, less researched genera of Heliconiini as well. Parapatry and sympatry are likely of importance in the ecological speciation of the butterflies, due to the dual role of wing patterns as both aposematic warnings under strong selection and essential cues for mating (Jiggins et al. 2001, Merrill et al. 2012). As incipient species diverge in their patterns by co-evolving with differently patterned co-mimics from multiple sympatric mimicry rings, assortative mating can lead to further genetic isolation of the lineages, enabling further divergence (Merrill et al. 2012). Conversely, the adaptive value of locally advantageous patterns facilitates adaptive introgression of pattern loci between species (Mallet et al 2007, Heliconius Genome Consortium 2012, Martin et al. 2013). Mimicry is thus an important factor in enhancing gene flow between species of *Heliconius* and *Eueides*, likely contributing to the observed levels of discordance between sampled markers. The seemingly unusual aspects of our study system, including introgression, hybridisation and speciation in sympatry, have all been only recently recognised as important evolutionary processes. We expect that reticulate signals in phylogenetic data will be increasingly important in future systematic studies and the Heliconiini represents a case study where the knowledge of specific microevolutionary processes can inform our understanding of cladogenesis at deeper timescales.

## Author contributions

Designed the study : KMK, NW and CDJ. Carried out the experiments, analysed the data and drafted the manuscript: KMK. Contributed novel materials and reagents: CDJ, JM, KD, NW and AN. All authors have read and approved the final version of the manuscript.

## Funding

KMK is funded by the Herchel Smith Trust, Emmanuel College and the Balfour Studentship from the Department of Zoology, Cambridge University. CDJ was supported by the Leverhulme Trust Leadership Grant. NW acknowledges funding from Kone Foundation. JM and KKD were funded through the BBSRC grant BB/G006903/1.

The authors declare no conflict of interest.

## Acknowledgments

Computational analyses were performed on the RAM—bio server at the School of Life Sciences, University of Cambridge, with extensive support from Jenny Barna. We are grateful to Brian Counterman, Laura Ferguson, Marcus Kronforst, Owen McMillan and Gilson Moreira for the permission to use unpublished Illumina data. We appreciate the specimens shared by Christian Brévignon, Luis Constantino, Frank Jiggins, Mathieu Joron and the curators at Harvard Museum of Comparative Zoology, McGuire Centre, Natural History Museum London and Naturhistoriches Museum Wien. We thank Nick Mundy for the permission to use vertebrate laboratory facilities. Jessica Leigh offered help with Concaterpillar. Rampal Etienne and Carlos Peña advised us on the diversification analysis. Members of the Butterfly Genetics Group, Rob Asher, Andrew Brower, Matthieu Joron, Nick Mundy, Albert Phillimore and Neil Rosser provided helpful comments on the results.

---

**Supplementary Figure S1.**
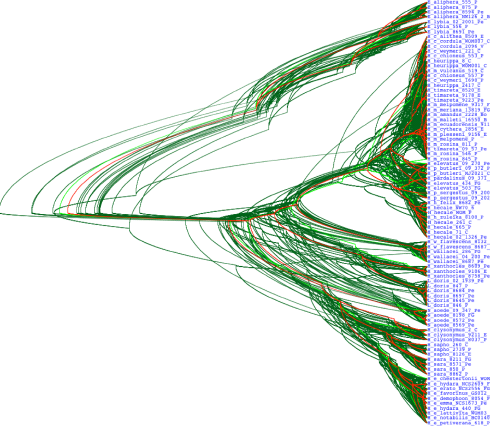
Gene tree estimates for well-sampled *Heliconius* species from the Bayesian multispecies coalescent analysis in *BEAST. Reticulation reaches the highest levels in the *Heliconius melpomene* clade.

**Supplementary Figure S2.**
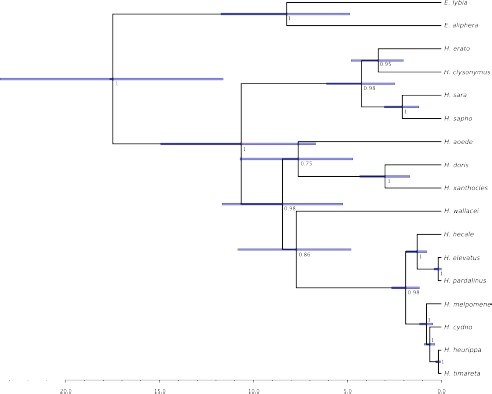
Species tree from the *BEAST analysis. The times of divergence were estimated using a relaxed molecular clock with the prior on the mean age of the root based on a dating of the Nymphalidae radiation (Wahlberg et al. 2009). Node bars show the 95% confidence intervals around the mean age. The scale axis indicates time in Ma.

**Supplementary Figure S3.**
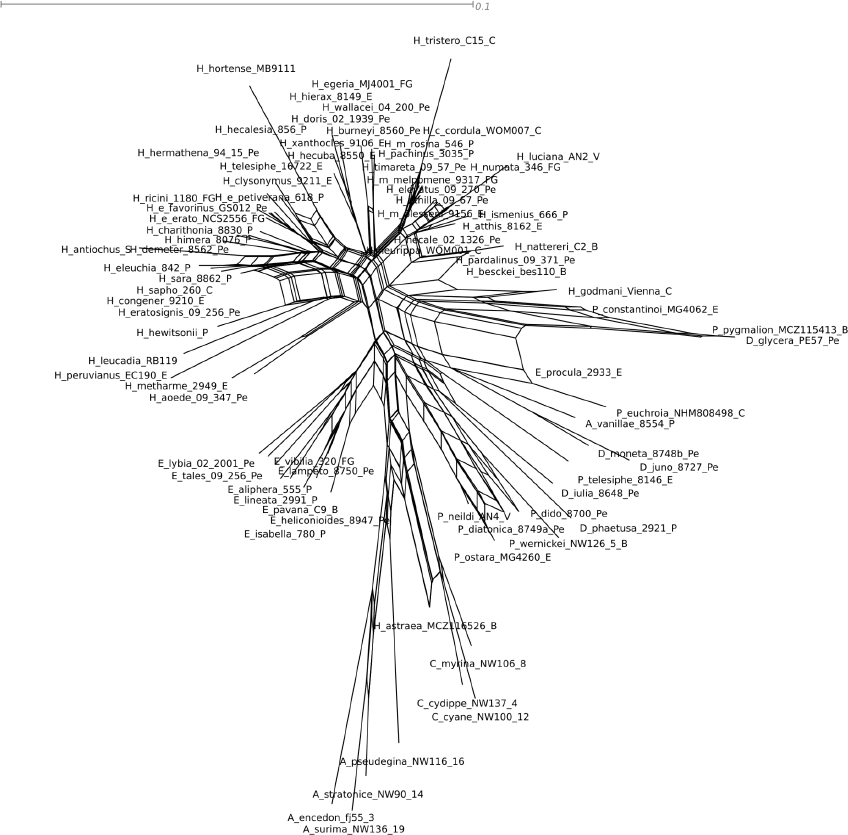
NeighborNet network of Heliconiini based on pairwise DNA distances computed under the F84 model of sequence evolution in SplitsTree v.4. Reticulation is especially pronounced at the vertices linking genera and subgenera of *Heliconius*. The scale bar is in units of average number of substitutions per site.

**Supplementary Figure S4.**
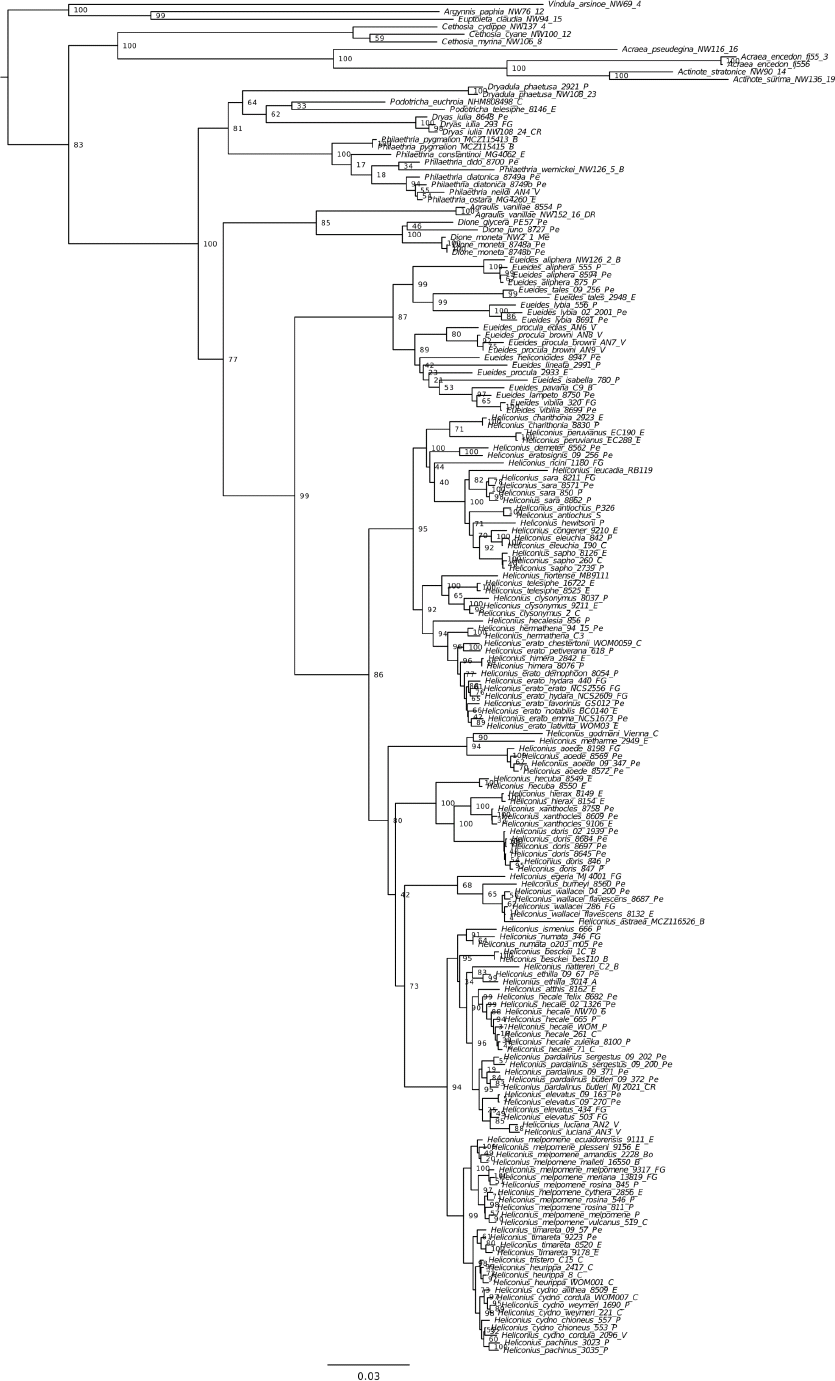
Maximum Likelihood (RAxML) phylogeny of Heliconiini butterflies based on two mitochondrial and 20 nuclear genes, estimated under the GTR+G model. Support values are estimated based on 1000 bootstrap replicates. All specimens are included in this analysis. The scale bar is in units of average number of substitutions per site.

**Supplementary Figure S5.**
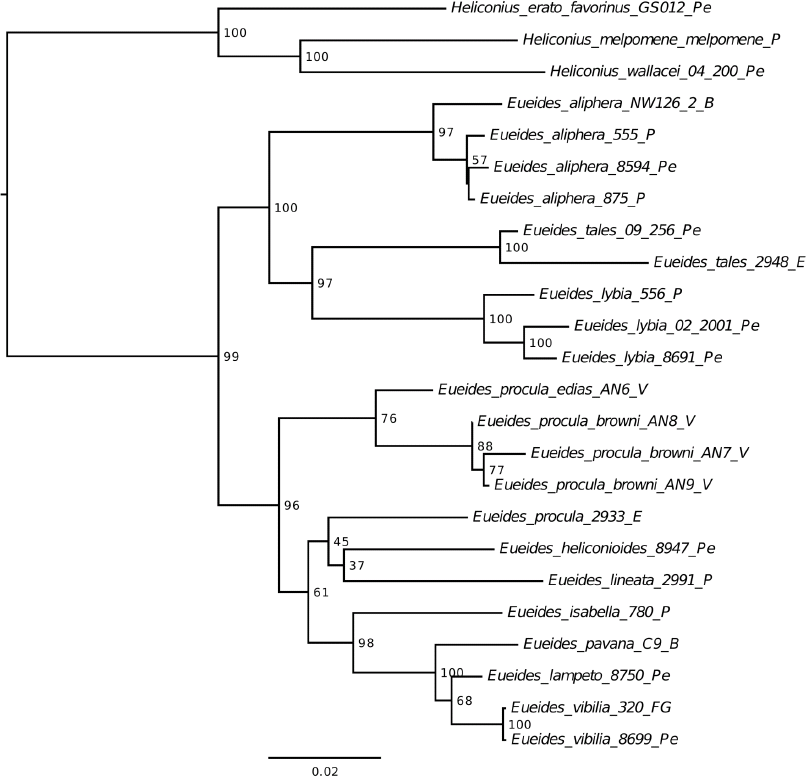
Maximum Likelihood (RAxML, GTR+G model) phylogeny of the genus *Eueides* based on 11 loci sampled in most of the species. Support values based on 1000 bootstrap repliactes, scale bar is in units of average number of substitutions per site.

**Supplementary Figure S6.**
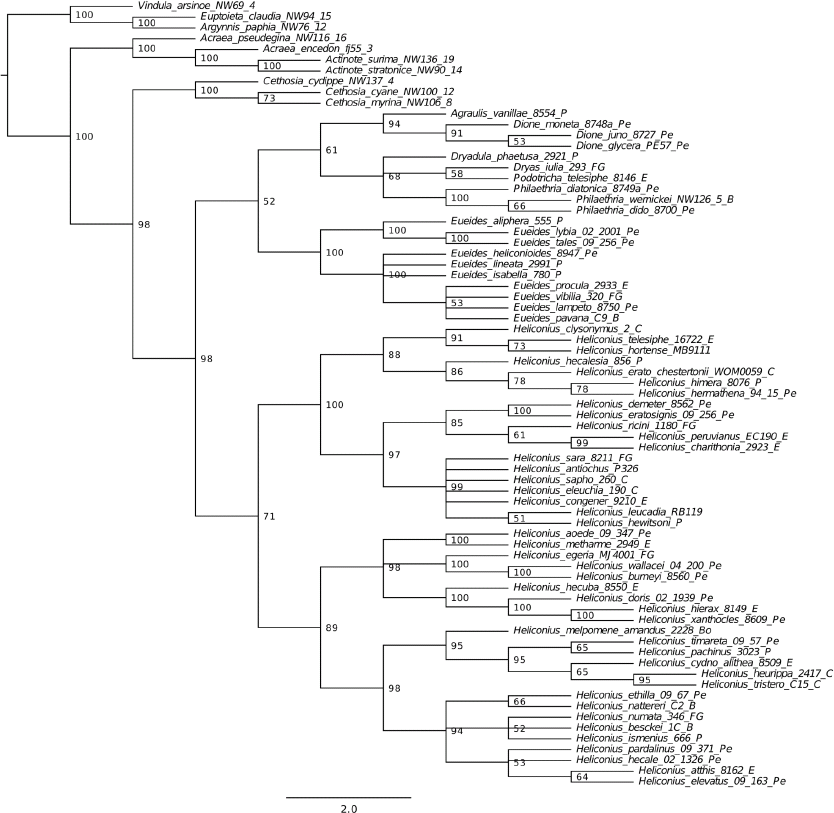
Minimise deep coalescences (MDC) phylogeny of Heliconiini: a 50% consensus of 100 bootstrap replicates of the set of 22 Bayesian gene trees.

